# *MetaStrainer*: Accurate reconstruction of bacterial strain genotypes from short-read metagenomic samples

**DOI:** 10.64898/2026.03.02.709061

**Authors:** Hazem Sharaf, Louis-Marie Bobay

## Abstract

**Summary:** Metagenomics provides broad insights from microbial communities, but more biological relevant phenotypes are attributed to subtle changes at the strain-level rather than species. Despite development of several tools using different algorithms, resolving individual strains from short-read pair-end sequencing data remains challenging. We developed *MetaStrainer*, a tool capable of reconstructing strain genotypes from metagenomic data. Compared with existing approaches, *MetaStrainer* substantially increases genotype accuracy, correctly identifies the number of strains, and accurately estimates their relative abundances. Accuracy of reconstructed genotypes is robust to choice of mapping reference.

**Availability and implementation:** MetaStrainer is implemented in Python 3. Source code and instructions are available on GitHub at https://www.github.com/lbobay/MetaStrainer and on Zenodo: https://doi.org/10.5281/zenodo.17872331

**Supplementary Information:** Supplementary data is available at Bioinformatics online.

## 1. Introduction

Metagenomic sequencing allows to explore the role that entire microbial communities play in their natural environment. Bacterial communities fulfil many key functions across virtually all environments. For example, studies of the human gut microbiome have revealed numerous associations between taxonomic composition and various health conditions [1]. However, despite substantial evidence supporting the role of the microbiome in human health, many studies still fail to identify specific microbial species associated with these conditions. This shortcoming has been attributed to the vast genomic variability that exists among strains of a single species [2,3]. Indeed, two conspecific strains can differ by >30% of their gene content and display distinct phenotypes, including metabolic functions, antibiotic resistance profiles and virulence factors. These findings highlight that the characterization of strains—rather than species—is essential for understanding the role that bacteria play in their environments.

Although metagenomic studies have provided many insights regarding microbial communities, analyzing these datasets still constitutes a major challenge. Multiple tools have been developed to assemble short reads *de novo* and generate Metagenome Assembled Genomes (MAGs) [4,5]. However, since metagenomic assemblers are largely unable to resolve individual genotypes in the presence of multiple related strains, the resulting MAGs capture only consensus species sequences and not the underlying strain genotypes. To address this issue, several bioinformatic tools have been developed to characterize strains within metagenomic samples [6]. These approaches fall into two categories: i) tools that *identify* strains using a set of gene markers and a database of Single Nucleotide Variant (SNV) markers (*e*.*g. MEGAN, StrainPhlAn*) and ii) tools that aim to *reconstruct* full strain genotypes using read mapping [7–9]. This latter category is more ambitious, and very few methods exist. Current tools like *StrainFinder* and *StrainFacts* rely on Expectation-Maximization (EM) algorithms to assign alleles to distinct genotypes based on allele frequencies [10,11]. However, these tools were originally developed to analyze multiple samples sharing identical strains and are not intended to reconstruct strain genotypes from samples with independent strain compositions.

More recently, *mixtureS* (also EM based), was released to reconstruct strain genotypes from independent samples [12]. Despite this progress, inferring strain genotypes from short read metagenomic data remains challenging.

We developed *MetaStrainer*, a tool that infers genome-wide strain genotypes from short read metagenomic data with high accuracy. *MetaStrainer* uses a framework distinct from previous approaches. First, it leverages paired-end read mapping to generate linkage groups of alleles. Second, it explores a wide range of possible strain distributions using a Markov Chain Monte Carlo (MCMC) search to identify the optimal strain composition and optimal strain phasing.

## 2. Methods

### 2.1 Implementation

*MetaStrainer* requires a user-provided genome as a mapping reference in GenBank format (gbff), and two paired-end metagenomic fastq files (forward and reverse reads). *MetaStrainer* can run with both complete or contig-level assemblies as a reference for read mapping. Because individual strains of bacteria often differ in gene content, the reference genome is first pre-processed to generate a database of individual genes against which the reads will be mapped.

Due to the nature of the read-mapping process and tapering coverage towards flanking ends, each reference gene is extended by adding flanking regions equal to the length of sequencing reads on both the 5’ and 3’ ends of each gene (*e*.*g*., 150bp on each side for 150bp reads, or less if the gene is near the end of a contig). By default, *MetaStrainer* assumes a read length of 150bp. *MetaStrainer* conducts the mapping of the reads against the reference database using Bowtie2 with *--very-sensitve --no-unal* options by default [13]. *MetaStrainer* interrupts sample processing when less than 60% of the mapping reference is covered as it typically indicates that the mapping reference is too distant to yield accurate genotypes (*i*.*e*., a different species).

Variant calling is then conducted on the reads, mapping the genes with a minimum base phred quality score of 30 and allele frequency between 1% and 99%. Flanking regions and gene positions with abnormally high or abnormally low coverage are excluded (customizable with a default at 1.5 standard deviations from the mean depth), and a minimum coverage of 20 reads per allele. is required. All allele variants detected within the same read or between paired reads are then linked into allele pairs (*i*.*e*., allele variants that are known to be physically associated). All the allele pairs are then joined together into linkage groups, which can span multiple genes.

In addition, non-reference alleles with 100% frequency are also tracked as they represent alleles that are not present in the mapping reference and are shared among the strains of the sample.

After constructing the linkage groups and computing their frequencies, *MetaStrainer*’s core algorithm is initiated by first assuming that three strains are present in the sample at equal frequencies. The three strains yield a hexamodal distribution where three peaks correspond to the frequencies of the alleles specific to strains 1, 2 and 3, respectively and three additional peaks correspond to the alleles shared by strains 1-2, 1-3 and 2-3, respectively. *MetaStrainer* then computes the distance of the frequency of each allele pair to the frequency of the closest peak, while doing the same for the complementary alleles. In the case where the complementary alleles are assigned to an incompatible peak (*e*.*g*., peak 1 and peak 1-3), the distance is arbitrarily set to 1. Finally, an overall score *S* is computed by summing the distances of all allele pairs. A MCMC search is then conducted using the MCMC sampler implemented in the Python package *emcee* [14]. The sampler is initialized with 500 random walkers following a Gaussian distribution, and with another sampler run from which random movements are drawn. The main algorithm iterates by drawing movements from these distributions, thereby adjusting the frequency of each strain genotype. After each movement, a slightly different hexamodal distribution of peaks is resampled using the MCMC sampler and the score *S* is recomputed on the new distribution. The mean frequency of the two first peaks are modified using two Gaussians of mean 0.005. The new distribution is accepted if the score *S* is lower than the previous step. Every 10 iterations, larger jumps are attempted by randomly selecting four types of jumps: the resampling of the means of the peaks is randomly increased by a factor 5, 10 or 100 or a new distribution is generated entirely. The same procedure of big jumps is initiated when 20 consecutive iterations have not identified a better distribution. The algorithm keeps on iterating until it converges towards a stable solution, which is defined by default as a series of 100 consecutive iterations where no better distributions are found.

In the final step, *MetaStrainer* reconstructs the genotypes from the strain distribution, the allele frequencies and the linkage groups. Using the allele-assigned hexamodal peak, *MetaStrainer* classifies the entailed allele to each strain based on a local confidence score which reflects how many times the allele is consistently paired in a relevant peak based on the hexamodal distributions. Alleles that are assigned to a different peak over 50% of the time relative to the other linked alleles in the linkage group are considered ambiguous and thus the allele is set to “N”. In order to avoid overestimating the number of strains due to small genotype differences, the reconstructed genotypes are then compared to one another based on a genome-wide average nucleotide identity. Nearly identical genotypes are collapsed into a single genotype (the most abundant one) using a customizable option with a default set at 99.5% genome-wide nucleotide identity.

### 2.2 Generating simulation datasets

Reads were simulated from *Gilliamella apicola*, a bacterial species that is part of the conserved core bee gut microbiome. The core genome of *Gilliamella* was obtained from [15]. The core genome was generated using CoreCruncher [16]. Reference strains were selected with an average nucleotide identity above 98% in their core genes (Table S2). Six genomes were selected, two of them were used as a mapping reference for aligning the Fastq reads, and the other four to create simulated populations (Table S1). Mapping strain A8 was selected as it had a 99.7% identity to A-1-24 strain, which is above the default strain delineation threshold of 99.5% we use to genotype strains.

Most species in microbiomes are usually dominated by a single strain, but several other less abundant strains may exist at much lower frequencies [9,17]. We thus generated samples composed of one to four strains at various relative frequencies (Table S3-5). CAMISIM was used to generate the simulated reads. For each simulated population, a configuration file with strain abundances was generated (Table S3-5). The simulation coverage for all strains in the populations is given Tables S6-9. Illumina built-in error profile *‘hi150’* was used to simulate 150bp reads sequenced with a HiSeq. A mean fragment size of 300bp and standard deviation of 30bp were used. All other settings were left at default. Prior to analysis, the raw simulated datasets were filtered from low quality reads using *Trimmomatic* v0.39 and the following parameters: LEADING:3 TRAILING:3 SLIDINGWINDOW:4:15 MINLEN:100 AVGQUAL:30 [18].

### 2.3 Evaluating accuracy of reconstructed genotypes

To evaluate the reconstruction genotype accuracy of each tool, a gene was assessed only if it was present in all simulated reference strains. (e.g. present in all three strains of the 3-strain simulations). All gene sequences of the reconstructed strains and reference strains were extracted and then aligned using muscle v5.1[19]. Reconstructed strains are matched to the original strain using the estimated frequency and the accuracy score. The accuracy score was calculated from all genome-wide variants in each, *i*.*e*. positions with different alleles between the reference strains and the mapping reference used. Positions including alignment gaps ‘-’, and ambiguous calls ‘N’ were excluded. Any position that did not include a predicted allele for a given tool was assigned the mapping reference corresponding allele and tested as such.

### 2.4 Comparing genotypes from different mapping references

To estimate the effect of the mapping reference on the tool’s accuracy (Fig. 1e,f), the orthologous genes of the two mapping references and the inferred genotypes were identified with *CoreCruncher*. The genes of all genotypes and references were then concatenated and aligned using *mafft* with the *‘–inputorder’* option [20]. The concatenated alignments for each tool were imported in R using the *ape* library version 5.6.2 [21]. A Hamming distance matrix was generated using the *dist*.*dna()* using *model=“raw”* and *pairwise*.*deletions=TRUE*. Nonmetric Multidimensional Scaling (NMDS) was done using the vegan package version 2.6.2 *metaMDS()* function with *k=3* [22].

**Figure 1:**
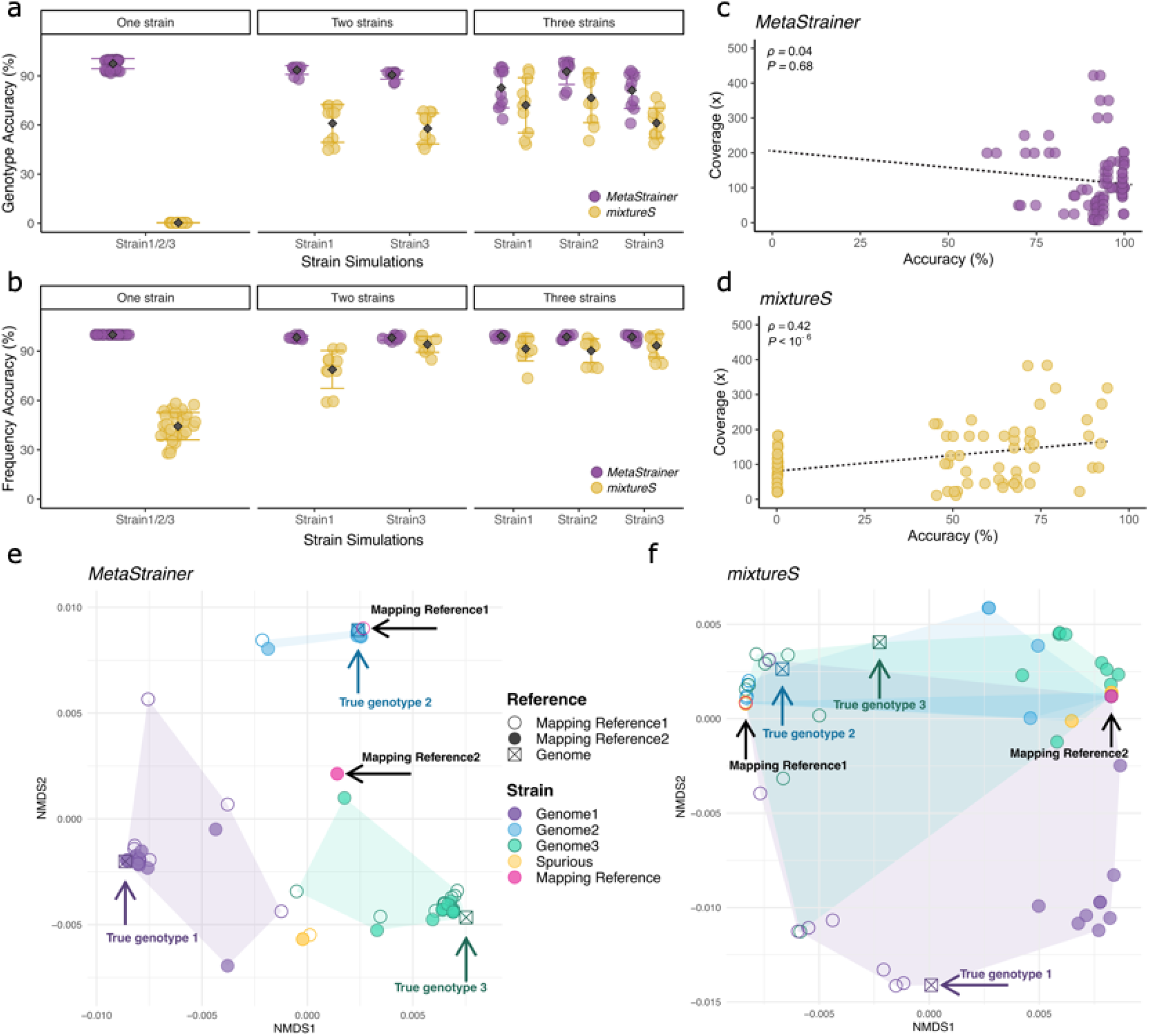
Performance of *MetaStrainer* and *mixtureS*. (a) Accuracy of genotype reconstructions. (b) Accuracy of the prediction of strain frequencies. (c-d) Impact of sequencing coverage on genotyping accuracy for *MetaStrainer* (c) *mixtureS* (d), respectively. (e-f) Non-metric multidimensional scaling (NMDS) ordinations measuring the relatedness of the reconstructed genotypes relative to the mapping references and the true genotypes for *MetaStrainer* (e) and *mixtureS* (f), respectively.

## 3. Results

We tested the performance of *MetaStrainer* using simulated metagenomes and compared its predictions to those obtained with *mixtureS* on the same metagenomes. Multiple metagenomic samples were simulated from full genomes of several strains of the bacteria *Gilliamella apicola* (Table S1) using CAMISIM [23]. For read mapping, we tested the tools using two reference genomes (strains A8 and N-22, Table S1). Each simulated population contained between one and three strains for a total of 56 simulations with various strain frequencies and sequencing depths (Table S3-5). In these simulations, *MetaStrainer* was able to infer the correct number of strains 53 out of 56 times (95%), whereas *mixtureS* predicted the correct number of strains only 4 out of 54 samples it processed (7%). The mean accuracy of strain frequency estimates for *MetaStrainer* was 99.1% and it was 71.3% for *mixtureS* (Figure 1b). We then evaluated genotype reconstruction accuracy, defined as the percentage of allelic variants correctly inferred.

*MetaStrainer* recovered 92.1% of the true genome variants compared to 39.3% for *mixtureS* (Figure 1a). As opposed to *mixtureS, MetaStrainer* does not genotype variants when the signal is too ambiguous (see Methods), and the unpredicted variants are designated as “N”. However, across the genotypes reconstructed with *MetaStrainer*, the number of uncalled alleles varied from only 0 to ∼200 (<0.86% of all variants) across samples.

We tested the impact of sequencing coverage on the ability of *MetaStrainer* to accurately reconstruct genotypes. We found no significant decrease in *MetaStrainer* accuracy relative to the sequencing coverage of each strain (Spearman’s *Rho*=0.04, *P*=0.69, Figure 1c), which was as low as 8X (5% frequency) for some strains. In contrast, *mixtureS* ‘s accuracy was affected by coverage depth (Spearman’s *Rho*=0.42, *P*<10^-6^, Figure 1d, Table S7).

We further analyzed how the choice of reference genome used for read mapping affected the accuracy of the genotype reconstruction. The two reference genomes used—strain A8 and strain N-22—share 98.8% identity across their core genome (Table S6). By design, strain A8 was chosen because it is nearly identical to one of the strains included in the simulated metagenomes. The PCoA plots in Figures 1d and 1e give an overview of the relationship of the genotypes reconstructed by *MetaStrainer* and *mixtureS* relative to the two mapping references and the true genomes used to simulate the three strains (*i*.*e*., what the reconstructed genotypes should be if the reconstructions were 100% accurate). *MetaStrainer* produced highly similar genotype and frequency estimates regardless of the reference genome used (*R*=0, *P*=0.4, ANOSIM, Figure 1D). In contrast, the genotypes reconstructed by *mixtureS* were strongly affected by the choice of the reference used for the mapping (R=0.94, *P*=0.001, ANOSIM, Figure 1E). Overall, *MetaStrainer* is highly robust to the choice of the reference genome used for the mapping.

Natural microbial communities are typically dominated by a single or a few strains per species, although additional strains can be present at very low frequencies [9,17]. The current implementation of *MetaStrainer* is limited to inferring a maximum of three strains. Alternative tools—such as *mixtureS*—do not have this limitation and are therefore expected to outperform *MetaStrainer* when samples are composed of four strains or more. To evaluate this, we tested both tools on simulated metagenomes composed of four strains (Table S1-5 and Figure S1).

*MetaStrainer* detected three strains (its maximum) in nine out of ten simulations, with a mean accuracy of 81.7%, whereas *mixtureS* detected at least four strains six times out of ten with a mean accuracy of 69.0%. *MetaStrainer* achieved more than 90% genotype accuracy for the dominant strain in six simulations (Figure S1c). Therefore, although *MetaStrainer* does not attempt to reconstruct all the strains present in the sample in these situations, those genotypes that are reconstructed are highly accurate.

Finally, because all strain-reconstruction algorithms rely on allele frequencies, their accuracy is expected to decline when strains are present at similar frequencies in a sample. We therefore tested *MetaStrainer* on the most challenging scenarios: strains present at equal frequencies (*e*.*g*., 1/3, 1/3, 1/3). As expected, the overall accuracy of the inferred genotypes was much lower (<70% of variants called accurately) in these extreme cases. Nevertheless, *MetaStrainer* still outperformed *mixtureS* in this situation (64.2% and 53.4% predicted variants, respectively, Figure S2).

## 4. Discussion and conclusion

Reconstructing the genotypes of larger numbers of strains remains challenging for *MetaStrainer* as well as for other approaches, including phasing or assembly-based methods, even those using long reads [24,25]. *MetaStrainer* and other algorithms are less accurate at reconstructing strain genotypes when i) many strains are present in a sample (≥4 strains) and ii) two or more strains occur at similar frequencies in a sample. However, in natural communities, most bacterial species are dominated by a single or very few strains [9,17], suggesting that methods focusing on a smaller—but more accurate—number of reconstructed strains are most relevant.

Newer tools have been developed using different approaches, such as phasing by minimum error correction of allele counts, and can more accurately identify the number of strains [24]. Therefore, we recommend using these tools prior to running *MetaStrainer* to estimate the number of expected strains in a sample. Overall, *MetaStrainer* is capable of reconstructing bacterial strain genotypes from metagenomic reads with high accuracy in most situations and outperforms the current tools for strain genotype reconstruction.

## Acknowledgements

We thank Faith Kennedy, Alessandro Oneto and Patrick Gallagher for testing the early versions of MetaStrainer. We also thank Kasie Raymann for input on the manuscript and the figures.

## Data availability

*S*ource code is available on GitHub at www.github.com/lbobay/MetaStrainer and on Zenodo: https://doi.org/10.5281/zenodo.17872331 Supplementary data is available at Bioinformatics online.

---

**Figure S1:**
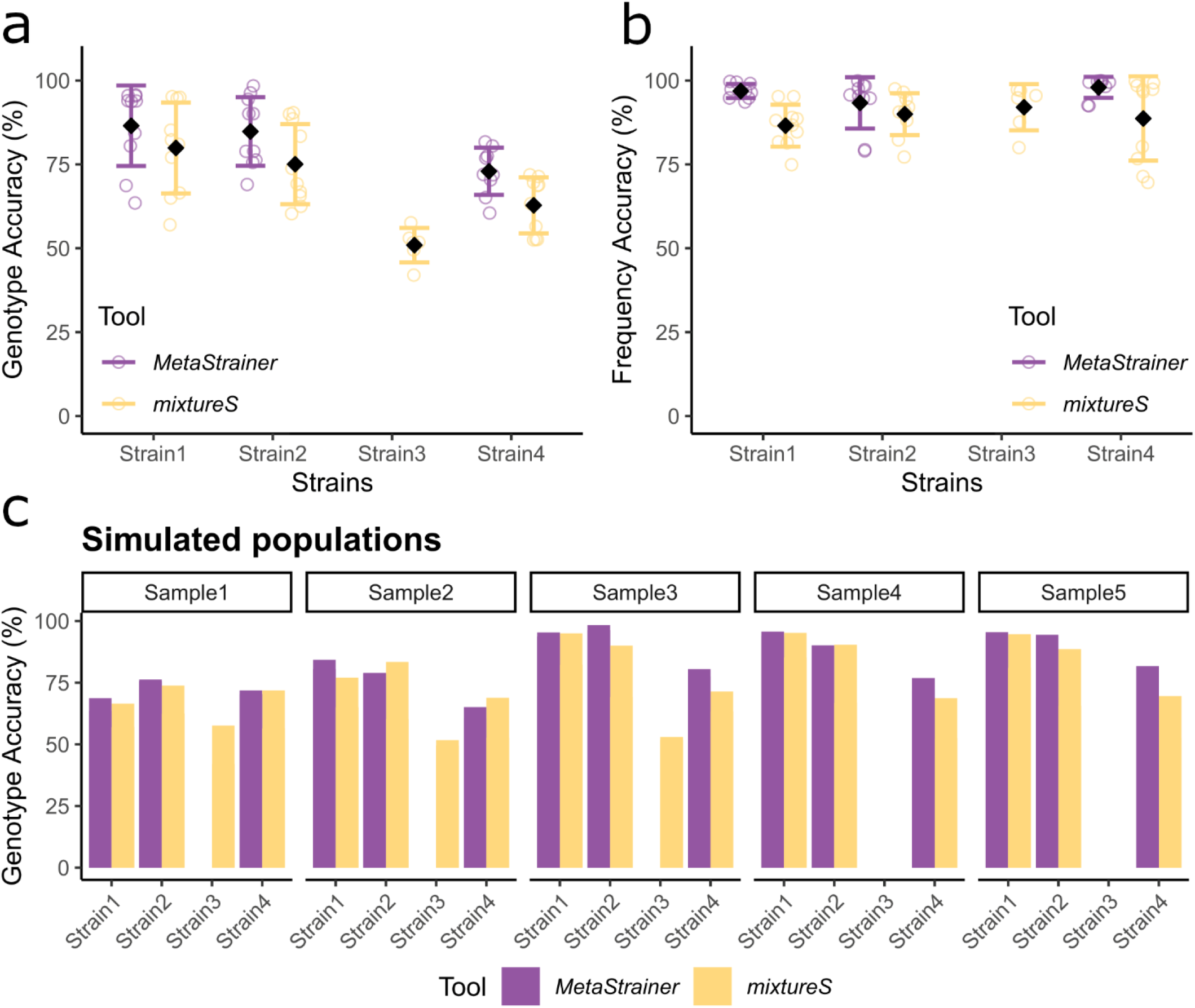
Genotype accuracy (a) and frequency accuracy (b) of *MetaStrainer* and *mixtureS* for samples simulated with four strains. Genotype accuracy (c) for each the populations simulated with four strains.

**Figure S2:**
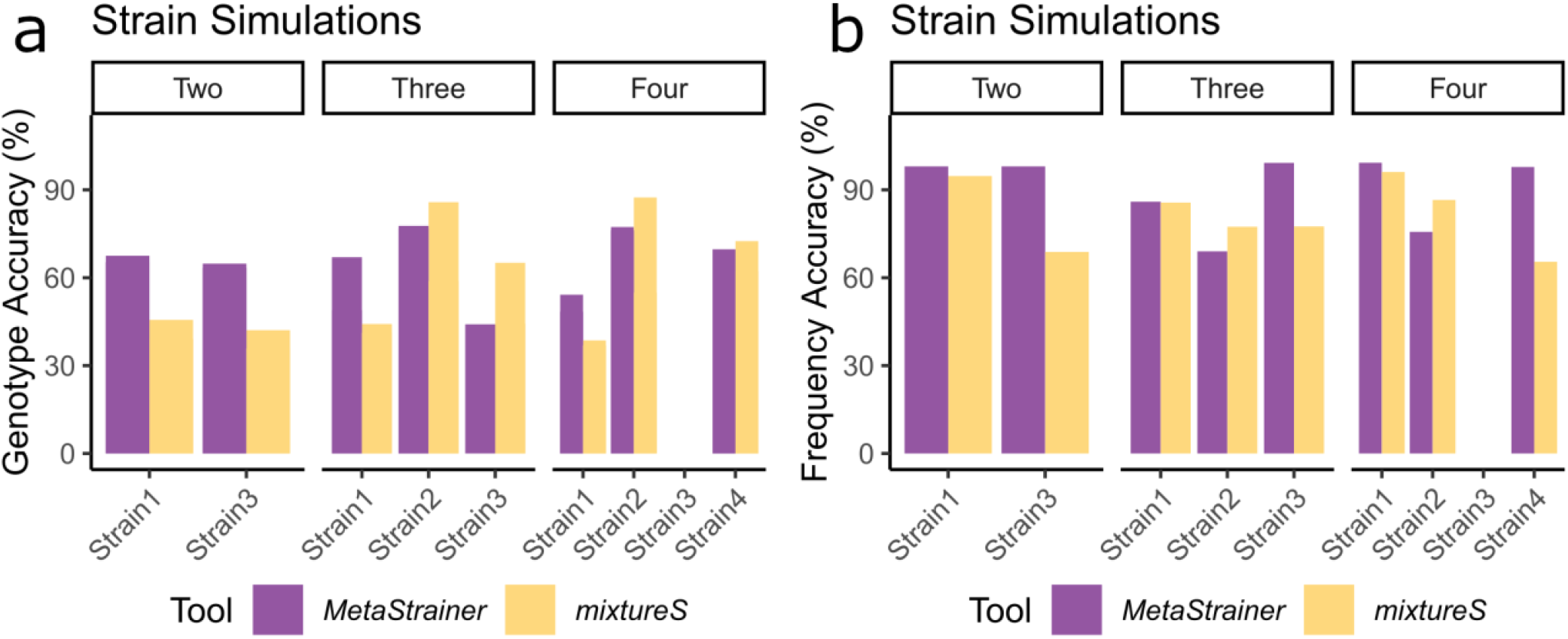
Genotype accuracy (a) and frequency accuracy (b) of *MetaStrainer* and *mixtureS* on samples simulated with strains at equal frequencies.

